# IBD stress impacts gut microbiome intra-species diversity

**DOI:** 10.64898/2026.07.02.736057

**Authors:** Chiara Mazzoni, Moran Yassour

## Abstract

Intra-species genomic variation results from diversity-generating processes and supplies the raw material for subsequent natural selection. Environmental stress can be regarded as the ultimate accelerator of these processes, especially for microorganisms, which can alter their DNA if presented with nutrient limitation, toxins, or pathogen attack. Chronic intestinal inflammation, as in inflammatory bowel diseases (IBD), may be regarded as prolonged environmental stress for gut commensal bacteria, bringing many enteric species down to undetectable levels.

However, it remains unclear how the microbes that survive the IBD gut environment actually respond to IBD stress, and whether their stress response may leave a transient or permanent signature in their genomes.

To investigate whether IBD stress induces and selects for certain genetic diversity, we performed metagenomic analyses on gut species in IBD patients and Controls. We focused on strain diversity within single individuals, which might be the result of more recent diversification processes under stress. We found measurable differences at the genome level between IBD and Controls, yet this was species-dependent. We then investigated gene-level diversity and found that certain functions were more likely to be enriched with either neutral divergence, functional divergence, or both. Functions that were enriched in IBD with both kinds of diversity were associated with motility and iron-scavenging, among others. These results may point towards functions that are under selection in the context of IBD stress, and could inform future mechanistic work, exploring previously unknown routes of bacterial diversification and adaptation to stress in the gut microbiome.

## Introduction

Host-associated microbiomes are generally considered more stable than free-living ones(1), as a host actively regulates its own internal bodily environment, promoting microbiome stability over time. Yet, host-associated microbiomes need to adapt to fluctuating conditions that affect the body, such as feeding cycles, seasonal shifts and climate changes(2–6).

In modern times, the human microbiome has been radically altered by the adoption of industrialized diets, sedentary lifestyle and use of medications(7–12). Many chronic diseases are globally on the rise, and many of these seem to be accompanied by changes in the human microbiome, particularly in the gut microbiome, the largest and most complex in the human body. Chronic intestinal inflammation, as in inflammatory bowel diseases (IBD)(13), alters profoundly both the host luminal environment and the resident gut microbiome composition(14). In order to clear invading organisms and defend the mucosal barrier, the host inflammatory response induces ample secretion of both biotic (defensins, lysozymes, secretory immunoglobulins) and abiotic (reactive oxygen and nitrogen species) antimicrobial agents(15–19). This antimicrobial response is meant to kill bacteria indiscriminately, but factors such as stress duration, intrinsic stress-tolerance and cell-to-cell interaction can affect whether certain strains are able to survive over others(20–23).

IBD is a complex and multifactorial disease that can be diagnosed as early as pediatric age, radically affecting the patient’s quality of life(24–27). Reduced gut microbial richness is a well-recognized hallmark of IBD, but beyond species-level shifts in the microbial composition, more research is needed to understand what characterizes the strains that tolerate IBD-driven stress in the gut(28, 29).

Tracking strain diversity has been largely used to determine the timeline and dynamics of single or multi-strain synthetic communities, while they acquire, accumulate and share genomic adaptations, both in stable and high stress conditions(30–33). Metagenomic sequencing allows the retrieval of genomes as they are present in their complex native communities(34), and with sufficient sequencing depth it can capture the presence of conspecific strains(35). To better characterize microbiome complexity and decipher microbial evolutionary dynamics, it is imperative to employ genomic approaches that take advantage of comprehensive genomic information. In human microbiome studies, metagenomics has been paired with multi-strain isolations and sequencing to resolve within-host diversification of phylogenetically related strains(36, 37). Employment of large cross-sectional metagenomic collections has been also used to infer species-specific evolutionary processes across hosts(38). These kinds of approaches have been used to identify and monitor individual strains over time, or describe scenarios of strain colonization, persistence and turnover, but mostly within healthy microbiomes, or specifically in the context of bacterial infections and fecal microbial transplants(39–41). More studies investigating microbiomes under stress are needed to understand the biological factors underlying bacterial survival to perturbation. This could potentially unlock avenues of clinical intervention in the context of antibiotic resistance, where exploiting dynamics of niche exclusion seems to hold promise(42).

Here, we employed a simple approach to track intra-specific genomic diversity across different gut commensals, comparing IBD patients to healthy individuals. Examining genomic positions that are polymorphic within a subject we evaluated the impact of IBD-stress on the gut microbiome. We employed nucleotide diversity, a metric commonly used in population genetics, as a measure of strain mixture within single subjects. We then expanded our analysis to evaluate the distribution and protein-changing potential of gene-level variants, distinguishing between nonsynonymous and synonymous ones. Analysing strain diversity, both at the genome and gene level, led us to define gene categories, which reflect gene-specific propensities to evolve under stress. To better understand mutational and selective processes in the context of stress, investigating metagenomic diversity could potentially inform microbiome remediation strategies in clinical and environmental settings.

## Results

### Gut commensals exhibit decreased or unchanged genome nucleotide diversity in IBD

In order to describe global strain-level diversity within individual hosts, we employed *nucleotide diversity*, a commonly used metric in population genetics (**Methods**). Nucleotide diversity is calculated as the average, per-position degree of polymorphism, over any nucleotide sequence length. It increases with the number of polymorphic positions and with the frequency of their minor alleles, peaking as alleles approach equal split. Coverage affects the number of positions that are included in estimating nucleotide diversity, as well as the number of minor alleles that can be confidently detected (**Methods**). In the context of this metagenomic analysis, nucleotide diversity is used as a proxy for within-host *strain mixture* or *richness*.

To compare nucleotide diversity (ND) levels between inflammatory bowel disease (IBD) patients and healthy individuals, we gathered metagenomic sequencing data from four different cohorts and three different countries, including gut microbiome samples from the Control group and both IBD subtypes (Crohn’s disease - CD, Ulcerative colitis - UC; **Methods**). We identified 20 gut commensals that could be robustly analysed across disease groups based on their prevalence across cohorts, and calculated their genome-level ND (**Supplementary Table 1, Methods**).

As displayed by the Control samples alone, ND estimates spanned different ranges across species, with *Blautia_A faecis* exhibiting the highest and *Agathobacter faecis* exhibiting the lowest median average ND (**Figure 1a**). Using per-species linear regression models, 13/20 species exhibited higher ND in Control compared to both IBD subtypes, while the remaining 7 species showed no difference, irrespective of existing differences in coverage breadth, coverage depth, relative abundance levels and different age group (**Figure 1b, Supplementary Figure 2a,c; Methods**).

**Figure 1:**
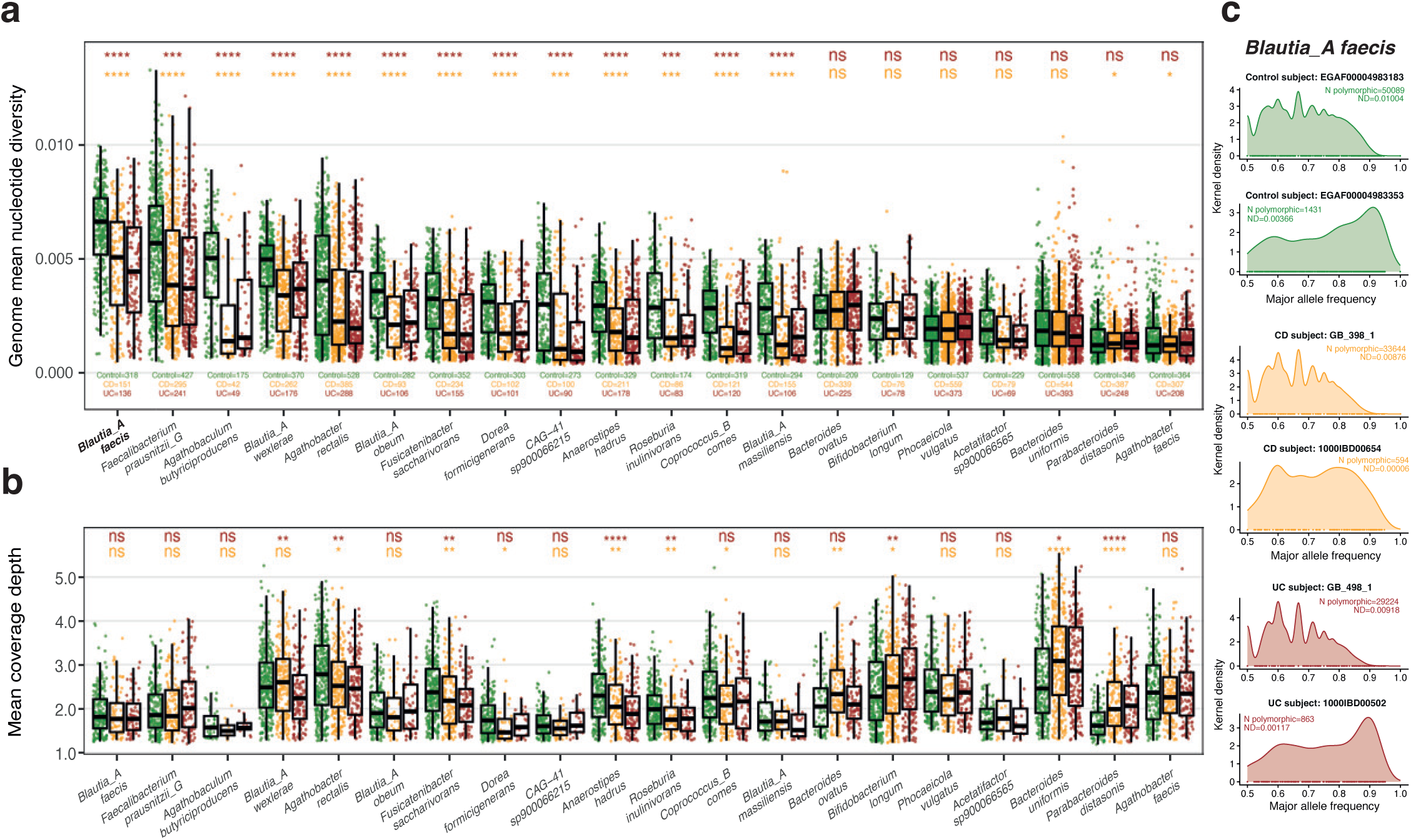
Genome mean nucleotide diversity differences across 20 gut commensals are independent of coverage levels. **a.** Genome mean nucleotide diversity per sample across 20 gut commensal species, comparing IBD groups (CD and UC - orange and red, respectively) to Control (green). Species ordered by decreasing Control group median. Group differences tested by species-specific linear-mixed modelling, adjusting for breadth and depth of coverage and age group (**Methods**). Statistical significance was corrected for multiple hypothesis testing using the Benjamini-Hochberg procedure. **b.** Mean coverage depth per sample, same order as in (a). Group differences were tested by Wilcoxon rank-sum test, corrected for multiple hypothesis testing using the Benjamini-Hochberg procedure. **c.** Major allele frequency density distribution across six different subjects (two per disease group) for the species *Blautia_A faecis*: annotations correspond to the total number of polymorphic positions along the genome (N polymorphic) and the genome mean nucleotide diversity (ND). Comparisons are statistically significant at alpha=0.05 and non-significant difference is reported as “ns” (p<0.05 : ‘*’, p<0.01 : ‘**’, p<0.001 : ‘***’, p<1e-04 : ‘****’).

Reliable ND estimation relies on accurate sequence alignment, hence we wondered whether systematic non-specific mapping across sequences of the same or different species could have potentially confounded our estimates. To account for this possibility and validate our results, we annotated species-specific marker genes within our species genomes (according to MetaPhlAn DB, **Methods**) and reran our models on this collection of genes (**Methods, Supplementary Figure 2b)**. When analyzing only the marker genes, we observed similar results to the full genome models, and the differences between IBD and Control groups were globally confirmed.

Both Control and IBD groups exhibited considerable within-group genome ND variation, and subjects in the same disease group could either display an enrichment of high frequency alleles or intermediate frequency alleles (**Figure 1c**). After establishing that our genome-level ND estimates were representative of the species core genome, it was clear that variance across samples reflected the multiple strains present within a subject, rather than possible differences in gene presence-absence. We selected two examples per disease group to illustrate how subjects could exhibit genome-level ND at the extremes of their group’s distribution, suggesting that health and IBD are not characterized by strictly high and low strain-level diversity respectively.

Using ND estimates as proxies for population-level strain diversity indicated that, on average, some species are subjected to prominent strain diversity loss in IBD, while others do not, and these changes recapitulate genetic diversity of core genes, independent from coverage levels.

### Only a few genes display increased nucleotide diversity in IBD

To explore whether there could be differences in nucleotide diversity (ND) levels across different regions of the genome, we next built a large set of linear regression models, one for each gene within a species genome (**Methods, Supplementary Figure 3**). We applied coverage-based and prevalence-based quality filtering of genes, and modelled disease differences, identifying gene-specific trends in ND levels. As a result, we categorized the genes based on whether their ND was invariant across study groups (*ND-invariant,* gray), or their ND was either lower or higher in IBD compared to Control (*ND-decreased,* blue; and *ND-increased,* orange, respectively; **Figure 2a, left-center**). Analogously to what was observed at the genome level, the same species that exhibited a prominent reduction in nucleotide diversity had a large fraction of their genes categorized as ND-decreased (such as *B. wexlerae* and *A. hadrus*). These species had ∼75% of their genes categorized as ND-decreased, with at most 47% gene overlap between CD and UC samples (intersection over union; **Figure 2a, right**). In a similar manner, species that exhibited no change in nucleotide diversity at the genome level, had most of the genes categorized as ND-invariant, with at most 82 individual genes categorized as ND-increased (in case of *B. uniformis*), which counted at most a 18% gene overlap between CD and UC. In order to validate the gene categorization that resulted from gene-level linear regression coefficients, we also calculated mean gene ND values (mean over subjects), distinguishing between Control and IBD, and calculated the IBD-over-Control ratio for ND-decreased and ND-increased genes (**Figure 2b**). Here, we confirmed that our gene categorization was valid, since ND-increased genes had higher IBD-to-Control ratio compared to ND-decreased genes (Wilcoxon rank-sum test, ND-decreased genes=586 [randomly sampled subset], ND-increased genes=45 [all]; **Methods**).

**Figure 2:**
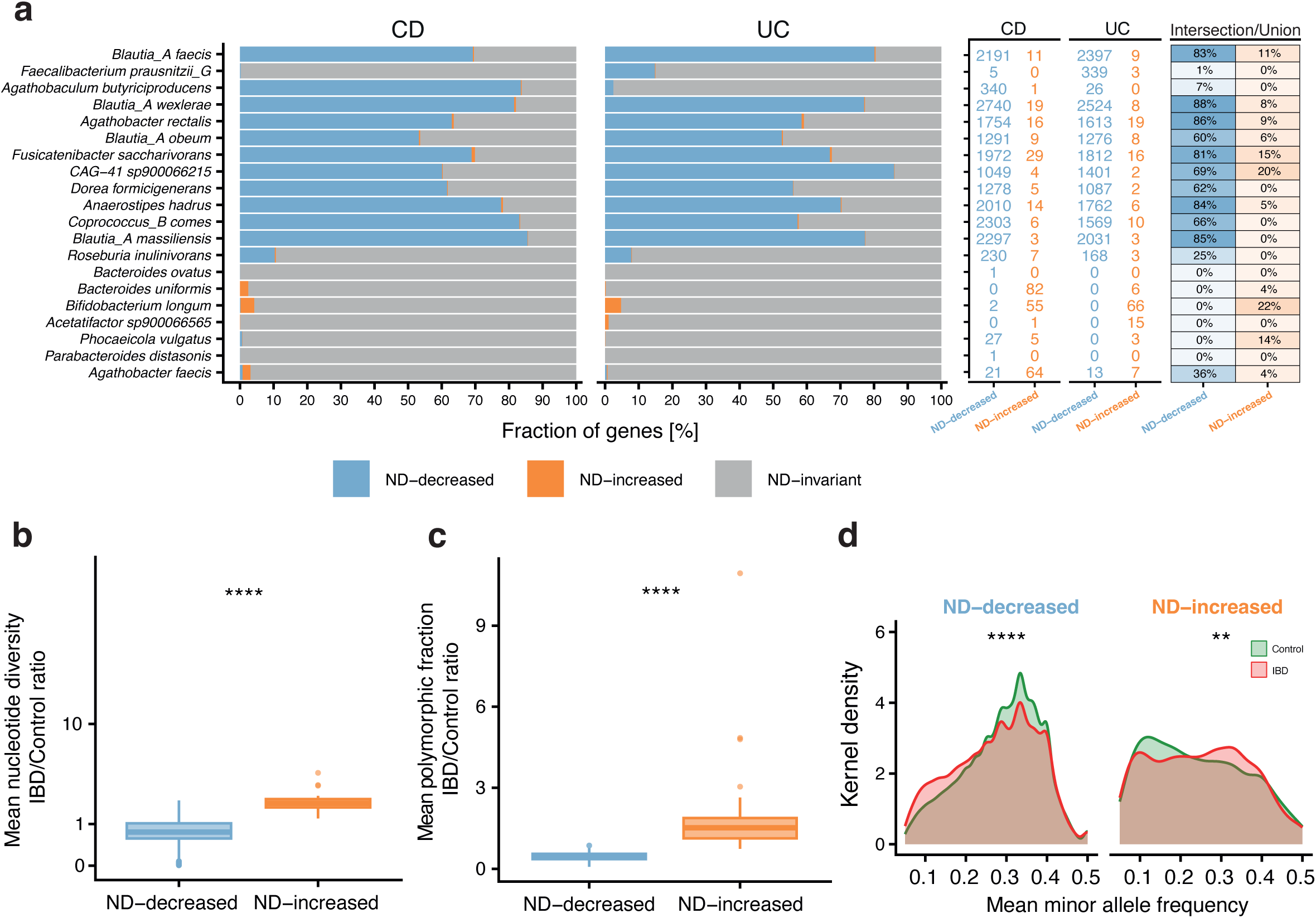
Nucleotide diversity enrichment in IBD defines 2 distinct groups of genes across gut commensals. **a.** ND-categorization overview based on linear regression coefficients (**Methods**):per-species fraction of genes colored by ND categories (ND-decreased - blue; ND-increased - orange; **left**), total number of genes in ND-decreased and ND-increased groups **(center)**, gene overlap percentage (intersection over union) between CD and UC subjects for ND-decreased and ND-increased groups **(right)**. **b.** IBD/Control ratio based on mean nucleotide diversity over subjects by disease group (IBD and Control). For this comparison ND-decreased genes were randomly selected (N=586), while all ND-increased genes were included (N=45). All selected genes were concordant in their ND category across IBD subtypes. **c.** IBD/Control ratio based on mean polymorphic fraction over subjects by disease group (IBD and Control), using the same selection of genes in (b). **d.** Mean minor allele frequency density distribution for ND-decreased genes and ND-increased genes as selected in (b,c). Differences in the distributions were tested using the Kolmogorov-Smirnov test. Comparisons are statistically significant at alpha=0.05 and non-significant difference is reported as “ns” (p<0.05 : ‘*’, p<0.01 : ‘**’, p<0.001 : ‘***’, p<1e-04 : ‘****’): p<0.05 : ‘*’, p<0.01 : ‘**’, p<0.001 : ‘***’, p<1e-04 : ‘****’.

Since nucleotide diversity estimates are affected by the number of polymorphic positions, as well as their allele frequencies, we investigated the contribution of each of these components to the ND levels measured (**Figure 2c-d**). We employed a similar strategy as before by using gene polymorphic fraction as the basis for calculating the IBD-to-Control ratio (**Figure 2c**). Also here, ND-increased genes had significantly higher IBD-to-Control ratio compared to ND-decreased genes (Wilcoxon rank-sum test; **Methods**). Finally, we tested the differential distribution of minor allele frequencies in ND-decreased genes and ND-increased genes, comparing IBD to Control subjects. Here, we observed significantly lower minor allele frequencies for ND-decreased genes in IBD subjects, while higher minor allele frequencies for ND-increased genes (Kolmogorov-Smirnov test, **Figure 2d**).

Focusing on the intra-species gene-level diversity, this analysis highlighted how individual genes, as units or subunits of cellular function, exhibited different levels of sequence diversity when comparing selected representatives of IBD- and Control-associated microbiomes.

### Nonsynonymous variants are enriched in IBD across species

To reason whether the observed trends in gene-level nucleotide diversity (ND) could be linked to adaptational processes happening in the IBD gut, we next investigated gene-level enrichment of nonsynonymous variants at polymorphic positions within subjects (**Methods, Supplementary Figure 4a**). We modelled genes’ relative accumulation of nonsynonymous and synonymous variants at polymorphic positions (*Nsyn-enrichment*) using a binomial generalized linear mixed-model on variant counts (**Methods**). For most of the 20 species, global Nsyn-enrichment was identified in both CD and UC, with few exceptions where only CD reached statistical significance (**Supplementary Figure 4b**). In the model, we included dNdS, which accounts for nonsynonymous enrichment at non-polymorphic positions, to approximate long-term genetic divergence between strains. Across species, dNdS was negatively correlated with Nsyn-enrichment, as shown by the negative model coefficients (**Supplementary Figure 4b**). Using model-derived gene-specific coefficients, we categorized genes in *Nsyn-decreased* (purple) and *Nsyn-increased* (yellow), based on whether they respectively exhibited a lower or higher Nsyn-enrichment in IBD compared to Control (**Figure 3a**, **Supplementary Figure 4c**). Some of the species that previously had no ND-decreased genes and few ND-increased genes, exhibited the largest fraction of Nsyn-increased genes (e.g. *A. faecis* and *A. sp900066565*; **Figure 3a, left**). CD and UC had a much more consistent overlap in Nsyn-increased genes (∼90%), with few species such as *B. ovatus*, *B. longum* and *P. distasonis* reaching only 51% or less overlap (intersection over union; **Figure 3a, right**).

**Figure 3:**
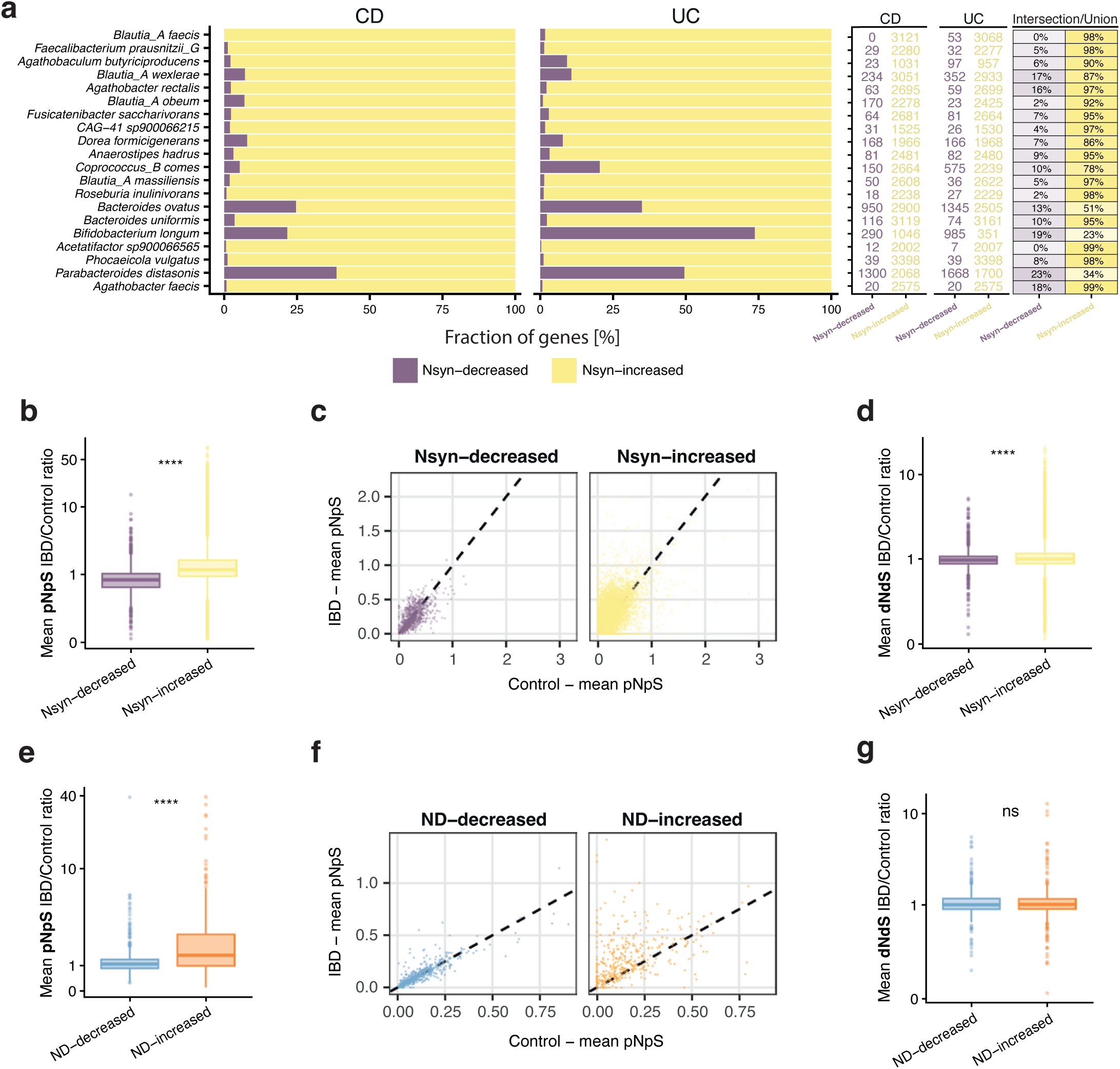
Nonsynonymous variants enrichment in IBD defines 2 distinct groups of genes across gut commensals. **a.** Nsyn-categorization overview based on linear regression coefficients (**Methods**): per-species fraction of genes colored by Nsyn categories (Nsyn-decreased - purple, Nsyn-increased - yellow; **left**), total number of genes in Nsyn-decreased and Nsyn-increased groups (**center**), gene overlap percentage (intersection over union) between CD and UC subjects for Nsyn-decreased and Nsyn-increased groups (**right**). **b.** IBD/Control ratio based on mean pNpS over subjects by disease group (IBD and Control). For this comparison all Nsyn-decreased genes (N=1283) and all Nsyn-increased genes (N=43263) were used. All selected genes were concordant in their ND category across IBD subtypes. **c.** Comparison between mean pNpS of Nsyn-decreased genes and Nsyn-increased genes in Control (x-axis) and IBD (y-axis) **d.** IBD/Control ratio based on mean dNdS over subjects by disease group (IBD and Control), using the same selection of genes in (b). **e.** IBD/Control ratio based on mean pNpS over subjects by disease group (IBD and Control) for ND-decreased (blue) and NS-increased genes (orange). For this comparison ND-decreased genes corresponded to the group randomly selected previously (N=586), while all ND-increased genes were included (N=45). All selected genes were concordant in their ND category across IBD subtypes. **f.** Comparison between mean pNpS of ND-decreased genes and ND-increased genes in Control (x-axis) and IBD (y-axis). **g.** IBD/Control ratio based on mean dNdS over subjects by disease group (IBD and Control). For this comparison ND-decreased genes and ND-increased genes corresponded to the group selected in (e). Comparisons are statistically significant at alpha=0.05 by Wilcoxon rank-sum test and non-significant difference is reported as “ns” (p<0.05 : ‘*’, p<0.01 : ‘**’, p<0.001 : ‘***’, p<1e-04 : ‘****’).

To validate that the Nsyn-enrichment identified by the model could be observed directly on genes using pNpS (nonsynonymous enrichment at polymorphic positions**, Methods**), we calculated IBD-to-Control ratios based on group-level mean pNpS, and tested gene category statistical difference (Nsyn-decreased=1283, Nsyn-increased=43263, **Figure 3b-c**). We confirmed Nsyn-increased genes had significantly higher mean pNpS IBD-to-Control ratio (Wilcoxon rank-sum test), as well as higher mean dNdS IBD-to-Control ratio (**Figure 3d**, Wilcoxon rank-sum test). Both pNpS- and dNdS-derived statistics confirmed that Nsyn-variants were specifically enriched in IBD across a subset of genes within the species’ genomes. Moreover, we also tested whether ND-categorized genes carried IBD Nsyn-enrichment, and only pNpS resulted to be differential between ND categories (**Figure 3e-g**, Wilcoxon rank-sum test).

These results showed that nonsynonymous polymorphisms were more prominent in IBD-associated strains compared to Control-associated ones and this was a common feature across the species investigated. The accumulation of nonsynonymous polymorphisms in IBD strains suggests specific mutational and selective processes happening in the IBD gut, potentially facilitating bacterial adaptation to environmental stress.

### Gene-level nucleotide diversity and nonsynonymous variants enrichment inform on gene evolutionary propensity under IBD stress

Thus far, we measured gene-level nucleotide diversity (ND) and gene-level nonsynonymous variant (Nsyn) enrichment. To offer a unified framework describing gene propensity to evolutionary change under IBD stress, we explored using ND and Nsyn model coefficients as axes of diversity, describing a two dimensional space, where genes can be localized in one of four gene categories: (1) *adaptive* with ND-increased & Nsyn-increased; (2) *neutrally-divergent* with ND-increased only; (3) *functionally-divergent* with Nsyn-increased only; and (4) *invariant* with ND-decreased & Nsyn-decreased (**Figure 4a**). As defined, adaptive genes exhibit higher genetic diversity in IBD along both ND and Nsyn axes, while neutrally-divergent and functionally-divergent genes exhibit higher genetic diversity along only one of the two axes. According to this framework, species exhibited differences in gene distribution and occupancy of the different categories, with visible differences between CD (top) and UC (bottom) patients (**Figure 4b, Supplementary Figure 5a**). Some species, such as *B. faecis* and *B. wexlerae* exhibited large variance along the ND axis, while *B. longum* and *P. vulgatus* exhibited only variance along the Nsyn axis. To investigate whether gene distribution across the four categories could be correlated with certain metabolic pathways or cellular functions, we computed functional enrichment propensities (logarithmic odds-ratios, log(OR)), grouping genes in KEGG pathways, BRITE hierarchical groups, EC numbers and KO terms (N functions with ≧5 genes=242; **Methods**). Specifically, we computed the odds-ratios of being categorized as *adaptive*, *functionally-divergent* or *neutrally-divergent* for each functional group (**Methods**). Odds-ratios for the adaptive category identified 62 functional groups, as either strongly enriched with- or depleted of diversity in IBD [*absolute log(OR)>=1*] (**Figure 4c, left**). For these functional groups, we highlighted both neutrally-divergent and functionally-divergent odds-ratios, together with species-specific gene contributions to the functional group (**Figure 4c, center-right**). Functional groups that had the highest adaptive odds-ratios (top of list) included bacterial motility (*Chemotaxis*, *Flagellar assembly*), iron binding and oxygen/redox sensing (*ferritin* and *hemerythrin,* respectively), secretion systems components (*Gram-positive* secretion component; *HlyD family secretion protein)*, outer membrane vesicle formation (*LemA protein*), sugar metabolism (*glucosidase*, *L−rhamnose mutarotase*, *neopullulanase),* and host-derived polysaccharide degradation *(Glycosaminoglycan degradation* and *unsaturated chondroitin disaccharide hydrolase).* Among the functional groups that were depleted of diversity (**Figure 4c, center-right,** bottom of list), we identified cofactor-associated functions (*copper homeostasis protein*, *cobalamin reductase*), enzymes involved in fatty acid metabolism (*acyl−CoA thioester hydrolase*, *oleate hydratase*), sulfur cycling and metabolism *(Sulfur relay systems, Glutathione metabolism*, *Sulfoquinovose metabolism*), and pore forming toxins *(Bacterial toxins* - *hemolysin III*). Finally, we also looked at functions that exhibited particularly large ND and Nsyn coefficients in individual species (**Supplementary Figure 5b**). Among these, we found several genes encoding for tRNA biogenesis proteins, transporters and factors involved in sensory signal-transduction cascades, including again proteins involved in chemotaxis and flagellar assembly.

**Figure 4:**
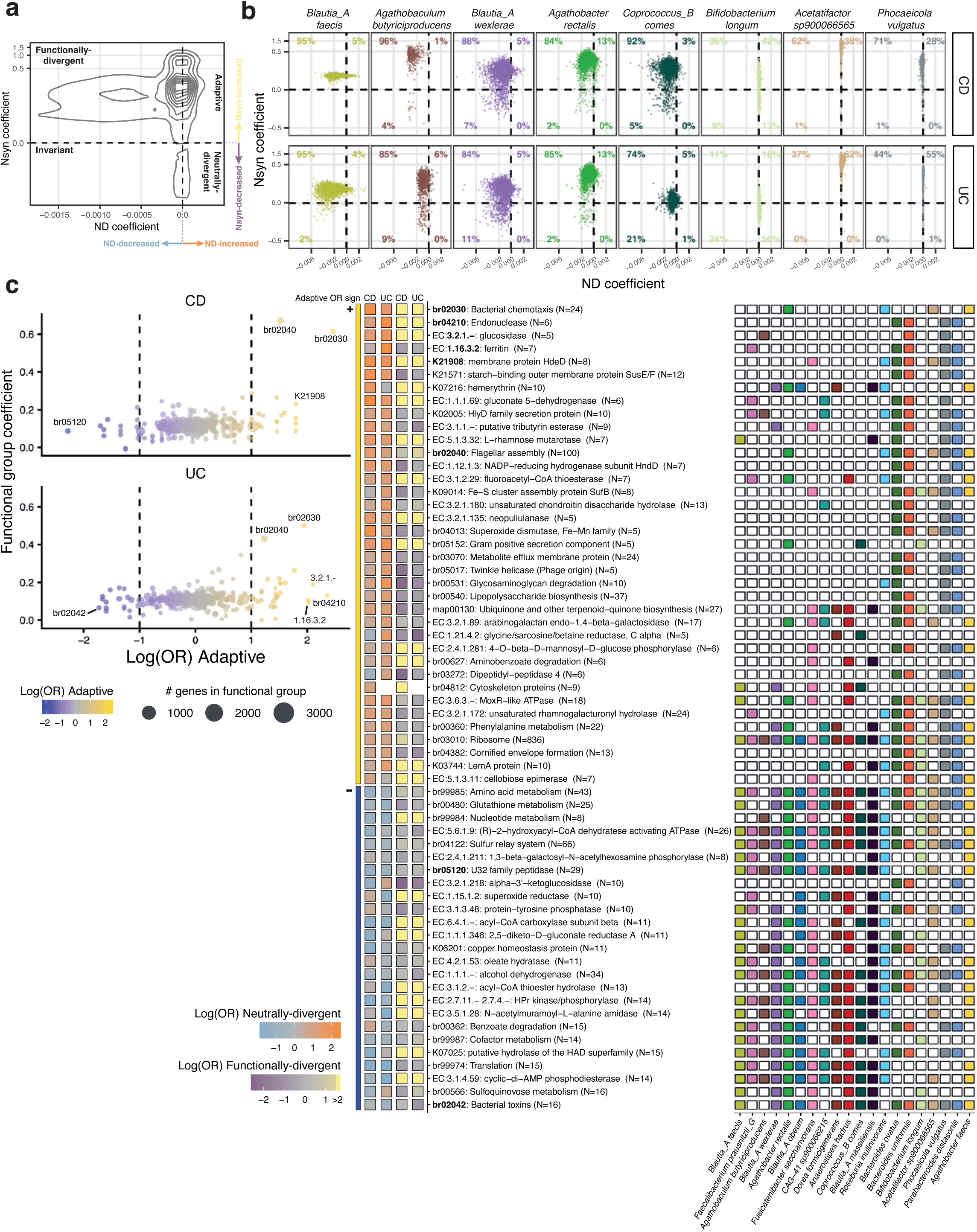
Gene-level nucleotide diversity and nonsynonymous variants enrichment suggest motility as the most adaptive function under I8D stress. **a.** Four categories of gene evolutionary propensity are described by nucleotide diversity (ND) coefficients (x-axis) and nonsynonymous variant enrichment (Nsyn) coefficients (y-axis). Contour lines describe the distribution of genes in these categories across species. **b.** Selected species-specific distribution of genes in the four categories of evolutionary propensity under IBD (CD on top, UC on the bottom). The gene fraction occupying each of the four categories is reported in percentage. **Supplementary** Figure 5a shows the rest of the species. **c.** (**left**) Functional group enrichment in the Adaptive category (ND coefficient>0 and Nsyn coefficient>0) across IBD subtypes, measured as logarithmic odds-ratios (x-axis) against the joint functional group-level coefficient, measured as the median of individual gene sums of ND coefficient and Nsyn coefficient (y-axis). Size of the dots corresponds to the number of genes and data points beyond the dashed lines exhibit strong functional enrichment. Some functions at the extremes are highlighted using their functional ID. (**center**) Functional groups with strong positive and negative enrichment in the Adaptive category (side sign bar) with their logarithmic odd-ratios for the other two evolutionary categories (squares, Neutrally-divergent and Functionally-divergent). Functions highlighted on the left are also reported here in bold. Labels report the functional identifier (either BRITE number, EC number or KO), a short definition and the number of genes in that category in brackets. (**right**) Species-specific gene contribution to the functional group: the panel reports with a colored cell whether a functional group includes a gene encoded by a certain species. The colors used for the species are the same as in (a).

These results showed that some genes, possibly related to bacterial stress avoidance (motility) or inter-specific competition (secretion systems), are likely to present signs of genetic diversification in IBD, suggesting higher propensity to evolve under stress, while other genes, involved in nutrient scavenging and cycling, might be devoid of genetic diversity for their essential roles in bacterial stress management.

## Discussion

Healthy microbiomes may support large and diverse populations of conspecific strains, which can usually stably colonize and diversify for years within the host(36, 40, 43–46). Factors that alter the diversity of the existing pool of strains in a community can be seen as ecological disturbances (e.g. bottlenecks), which filter out poorly adapted strains before allowing for recolonization events(47–50).

IBD acts as a strong and prolonged ecological filter, with escalating bouts of gut inflammation causing drastic remodeling of the microbial community within the gut(51). As we showed in the first section, more than half of the species tested in our analyses exhibited signs of reduced strain richness in IBD, while the remaining few showed no change. On one hand, reduced strain richness in IBD could result from the lack of either strain retention or recolonization processes, but in both cases the IBD gut exhibits overall lower ecosystem carrying capacity, lacking open and stable niches that can be occupied, potentially due to mucosal inflammation(52). On the other hand, the lack of change in the diversity of some species across disease groups is surprising, given that drastic compositional changes are observed (**Supplementary Figure 2c**). For these species, one possible interpretation could be that already at homeostasis, a few strains tend to dominate, and the IBD gut has the potential to accommodate all of them(40, 53). Alternatively, these species might be characterized by intrinsic capacity to survive and adapt to the IBD stress environment. The species that exhibited no change in our analysis, such as *P. vulgatus*, *B. uniformis* and *B. longum*, are not opportunistic facultative anaerobes, such as the ones belonging to the *Enterobacteriaceae* family, which easily adapt to the more aerobic gut environment driven by inflammation(54, 55). These species are prevalent and at times highly abundant human gut species, suggesting that high prevalence and abundance might indeed be signs of high resistance to ecological stress.

Resilience to IBD stress could be encoded in a species metabolic capacity, in particular by encoding complete metabolic modules enabling for the biosynthesis or assembly of essential nutrients and cofactors, which are often turned into rare commodities under gut inflammatory conditions(29, 56–58). Thus, our analysis focused heavily on genetic diversity at the individual gene level, where we identified a specific gene subset across species that exhibited an increase of nucleotide diversity in IBD, in contrast to the rest of the genome (ND-increased genes). This suggested that a gene-level diversity analysis has the potential to identify gene functions that tend to accumulate polymorphisms, and that the retention of these polymorphisms might be explained by an acquired survival advantage in the context of IBD stress.

Additionally, we measured the overall enrichment of nonsynonymous diversity at the gene level and identified gene-specific enrichments unique to IBD strains. These analyses suggest that overall IBD-associated strains seem to undergo a specific selection process, where functional innovations (nonsynonymous variants) are selected for. Also in this case, some species exhibited a more prominent nonsynonymous variant enrichment than others, not necessarily overlapping the nucleotide diversity enrichment. Assuming IBD stress exerts high purifying selection pressure on the gut microbiome, the identified functional variants in IBD-associated strains could be likely the result of recent adaptive diversification, rather than relaxed purifying selection(59). The trigger to these nonsynonymous mutations could be the exposure to oxidative stress (ROS and NOS), which triggers DNA damage(60–62). When DNA damage occurs, such as the oxidation of a guanine (8-oxo-G), stalled DNA replication triggers recruitment of an error-prone DNA polymerase, which often incorporates the wrong nucleotide, increasing the likelihood for nonsynonymous codon change. The process of stress-induced mutagenesis is very prominent in the context of a variety of environmental stresses, including antibiotic stress(63, 64). Another trigger for the biased incorporation of nucleotides under stress could be nutrient limitation. It has been described that nutrient limitation was the basis of evolutionary variants among marine bacterial genomes, specifically under limited availability of nitrogen(65). In IBD stress, many nutrients are limited, though whether this kind of nutrient limitation is responsible for specific evolutionary innovations in IBD remains unknown.

Finally, we focused on combining all the evidence in one unified framework. We defined four gene categories describing differential propensity to evolutionary change under stress, and identified functions that were enriched in overall nucleotide diversity and nonsynonymous variants in IBD (*adaptive* functions). We highlighted motility, secretion system components, oxidative stress management proteins and many others that open interesting avenues for future mechanistic work. Specifically, the gene group associated with flagellar assembly was one of the biggest functional groups enriched in the adaptive category. Some studies have highlighted that mutated flagellar components may induce cell aggregation, allow for better host colonization, or possibly reduce intracellular ROS production and mutagenesis(66–70). In the context IBD stress in the gut, further experimental work will be needed to elucidate whether increased genetic diversity in flagellar components positively or negatively impact survival in the gut by optimizing or inactivating bacterial motility.

We also identified diversity enrichment in proteins associated with housekeeping functions. Among them, we found ribosomal proteins, which are commonly used as universal marker genes to infer phylogenetic relationships between distantly related bacteria(71, 72). Recent work has performed a large-scale phylogenetic analysis using a set of universal marker genes (including ribosomal proteins and an RNA polymerase subunit) and identified IBD-associated strains that were later confirmed to be likely IBD-adapted, based on the metabolic functions that they encoded(29). This work suggested that IBD-associated strains might carry an evolutionary signature in their universal marker genes, and this signature might derive from stress exposure. Other *in vitro* studies on *E. coli* confirm that, in response to long-term starvation, strains reproducibly acquire mutations in RNA polymerase subunits (rpoB and rpoC), to the extent that coexisting strains can be tracked over time by means of RNA Pol subunit barcoding(73–75). Collectively, these findings point towards the occurrence of global genetic remodeling in bacteria under stress, and understanding the nature of these genetic changes might further uncover unknown strategies of bacterial tolerance and interaction with the environment.

This work provides robust findings on the group-level relationship between disease and strain diversity, but also ignores sources of heterogeneity within disease groups, such as severity level and disease duration. These factors might be investigated in future work to elucidate whether remissive or active disease states are associated with different levels of strain diversity and whether disease duration could be associated with different kinds of genetic diversity.

Furthermore, longitudinal sampling of individual patients was only used to maximize species retrieval, but in future work this could be an interesting direction to investigate temporal dynamics of the genetic diversity observed cross-sectionally.

In the past decade, the study of host- and environment-associated microbiomes has uncovered how microbial communities can be used as dynamic barometers of organismal and ecosystemic health. This work highlights the potential of using intra-species gene-level diversity as a sensor for environmental stress, and suggests the presence of stress-derived evolutionary signatures in the genomes of commensal species in the IBD gut.

## Materials and Methods

### Metagenomic data cohorts

We compiled a collection of gut metagenomic sequencing data amounting to 2397 samples from the following cohorts: (1) the SZ pediatric cohort including 134 samples (Control=29,CD=61,UC=44), available at BioProject PRJNA1265906; (2) the LifeLines-DEEP_followup adult cohort including 337 samples from healthy individuals available under Data Access Agreement with the UMCG Department of Genetics to access EGAD00001006959 data repository; (3) the 1000IBD adult cohort including 352 samples (CD=203,UC=149) available under Data Transfer Agreement with the University Medical Center Groningen and Prof. Dr. R.K. Weersma to access EGAD00001003991 and EGAD00001004194 data repositories; the HMP2 mixed cohort including 1574 samples (Control=424,CD=714,UC=436) available at https://ibdmdb.org.

### Selection of the species

We identified species suitable for our strain-level metagenomic analysis by calculating species prevalence across the cohorts described above. We used Metaphlan4(76) and Metaphlan DB v. mpa_vJan21_CHOCOPhlAnSGB_202103 to taxonomically profile the samples, then we filtered out species instances with relative abundance < 0.01% and selected species whose prevalence reached at least 50% in each study group (Control, CD, UC). Since SZ and HMP2 cohorts include multiple samples from the same individuals, we calculated prevalence by considering any one sample with the species present per subject. From here we retrieved a list of 20 species whose prevalence and median relative abundance is reported in **Supplementary Table 1**.

### Cross-database mapping

The taxonomic identifiers for the 20 species selected were cross-mapped to the UHGGv1 database (taxonomy derived from GTDB release89) to identify corresponding representative genomes. We used the mapping file available with the Metaphlan DB (`mpa_vJan21_CHOCOPhlAnSGB_202103_SGB2GTDB.tsv`), then checked that the species representatives reported in GTDB release89 did not change till release202, ensuring that these representative genomes were stably identified as such for these species. Only one within our 20 species (*Phocaeicola vulgatus* SGB1814) was not present in GTDB release89, but it was introduced in release202 with GCA_964248265.1 as a representative genome. Hence, we retrieved GCA_964248265.1 from NCBI and renamed it GUT_964248265 to uniform it with other UHGGv1 representatives. The list of the 20 species with their SGB identifiers, GTDB naming and UHGGv1 identifiers is available in **Supplementary Table 2**.

### Strain-level metrics from metagenomic reads with inStrain

#### inStrain database

We used inStrain(77) to retrieve strain-level metrics directly from metagenomic reads. We first downloaded the UHGGv1 full set of representative genomes, identified coding regions with Prodigal(78) and built a bowtie2(79) database including genome GCA_964248265.1 as a representative for *Phocaeicola vulgatus.* Then we mapped all samples against the UHGGv1 DB using bowtie2 and samtools(80) to sort and index the mapping files. Then we ran inStrain with `--database_mode` which sets some parameters that are appropriate for mapping metagenomic reads to an external DB of genomes. This mostly ensures that the genome ANI threshold used is (1) not too stringent to allow for sample-specific strains to be mapped to their representative genome in the database, and (2) not too relaxed to ensure accurate, species-specific sequence mapping. Additionally we used different read pairing parameters based on whether the metagenomic samples were paired-end or single-end: for the single-end read samples (SZ cohort) we used --non_discordant, while for all the other paired-end read samples we used --paired_only (default).

inStrain by default filtered out samples that had less than 1x coverage depth for a species of interest, so the number of samples that were positive for at least one of the 20 species selected was 1435.

#### Coverage breadth filtering and the B. longum case

We gathered all inStrain genome-level outputs and filtered out instances where a species of interest did not reach a minimum coverage breadth threshold. The recommended minimum coverage breadth by inStrain documentation is 0.5, but we noticed that, applying this threshold, one of the 20 species selected, *Bifidobacterium longum*, was underrepresented in our dataset compared to our prevalence expectations. The UHGGv1 database indeed includes a *B. longum subsp. infantis* representative genome and this likely resulted in the lower coverage breadth observed for *B. longum* in our data, which likely belongs to *B. longum subsp. longum*. In **Supplementary Figure 1a** it is visible how all instances of *B. longum* genomes are indeed right below the recommended coverage breadth threshold, so we decided to use 0.45 as the coverage breadth threshold specific to *B. longum*. Additionally, we filtered out species instances where the genome had at least a coverage breadth equal to 0.2 at a minimum coverage depth equal to 5x (`breadth_minCov >=0.2`, `minCov=5x`, default). Applying these coverage filtering thresholds, 1408 samples were positive for at least one species of interest.

#### Coverage depth and nucleotide diversity

Among the metrics provided by inStrain, we started analysing nucleotide diversity (ND). inStrain calculates ND according to its original implementation(81) and by default only using positions that reach a minimum coverage depth threshold (`minCov=5x`). ND is provided both at the genome and gene level, but to have better control on ND estimation accuracy we decided to use only gene-level ND estimates for genes that had breadth_minCov >=0.2.

### Linear regression modelling

#### Genome-level nucleotide diversity

To obtain genome-level nucleotide diversity averages, we calculated the mean nucleotide diversity across genes previously filtered by `breadth_minCov`, adjusting nucleotide diversity by the product of `breadth_minCov` and gene length, to account for nucleotide diversity estimates calculated from poorly covered genes, without raising the general threshold for gene inclusion. We did the same for calculating the genome-level mean for coverage depth and breadth and compared the unweighted and weighted genome-level nucleotide diversity means as shown in **Supplementary Figure 1b**. Then we used these weighted metrics to run a linear regression model estimating the effect of disease on genome-level nucleotide diversity per species, adjusting for coverage depth and breadth and age group, as follows: lm(`calc_genome_metric ∼ Phenotype + log(calc_genome_coverage) + calc_genome_breadth_minCov + cohort, weights = breadth_minCov* gene_length)`.

Results from this set of models are reported in **Supplementary Table 3** and **Supplementary Figure 2a**. As input to the linear regression model, we used only cross-sectional data, so for subjects with more than one sample available, we selected the sample with the highest genome-level mean `breadth_minCov`. Finally, we collected model results for all 20 species, used the lmerTest (v.3.2-1) R package to retrieve statistical significance for the model coefficients and applied the Benjamini-Hochberg procedure to account for multiple hypothesis testing.

#### Marker gene-level nucleotide diversity

To evaluate whether the results from the genome-level nucleotide diversity means could reflect variability across genes included in the mean rather than the whole genome, we decided to rerun the linear regression model using nucleotide diversity means coming from core genes only. To identify genes that could represent the core genome, we searched for the Metaphlan DB species-specific marker genes in our set of genes. For this, we mapped the 20 species representative genomes against the Metaphlan DB (v. mpa_vJan21_CHOCOPhlAnSGB_202103) using BLAST(82) and then selected for the best alignments using their bitscore. Out of 2388 total markers associated with the 20 species, we confidently identified 2296. Then we calculated genome-level mean as described previously and ran the linear regression model as before. Results from this set of models are reported in **Supplementary Table 4** and **Supplementary Figure 2b**.

#### Gene-level nucleotide diversity

To obtain gene-level disease effects on nucleotide diversity we ran a large set of gene-level linear regression models, since parametrization for the full set of genes in the previous model set did not manage to converge, even with the introduction of different optimizers. For this reason, we ran a linear regression model per gene and then collectively applied bayesian shrinkage using the R package ashr (2.2-63) to reduce overinflated coefficients. Genes that did not reach per study group a prevalence>0.2 and were present in a minimum number of subjects of 10 were not used to build a model. The models were run as follows using stats R package (v. 3.6.2): `lm(gene_metric ∼ Phenotype + log(coverage_scaled) + breadth_minCov + cohort, weights = breadth_minCov* gene_length)`.

According to this procedure, genes that had a Local False Sign Rate (lfsr) < 0.05, derived from shrinkage were considered to have a statistically significant coefficient and if the genes had coefficient<0 were termed as “ND-decreased”, while if the coefficient>0 they were termed as “ND-increased”. To confirm that the gene categorization that resulted from the linear regression model corresponded to our expectations in (1) mean nucleotide diversity, (2) polymorphic fraction and (3) minor allele frequency, we selected the genes that were categorized concordantly across CD and UC, for both ND-decreased and ND-increased genes, and then calculated the IBD-to-Control ratios based on the three different metrics mentioned and tested statistical difference (ND-decreased genes=586 [randomly-sampled subset], ND-increased genes=45 [all])

#### Gene-level nonsynonymous variants enrichment

To model the relative gene-level accumulation of nonsynonymous and synonymous variants, we decided to use raw variant counts (`SNV_N_count` and `SNV_S_count` in inStrain, respectively) in a binomial generalized linear mixed-effects model using glmmTMB function from glmmTMB (v. 1.1.13) R package, as follows: `SNV_N_count, SNV_S_count ∼ Phenotype + dNdS_substitutions + log_cov_scaled + breadth_minCov + (1 + Phenotype || gene), family = binomial(), control = glmmTMBControl (optimizer = nlminb).` Thanks to the binomial structure, the model interprets data from genes with few variants as less reliable, and with gene-specific random effects, as in `(1+Phenotype|gene),` we are able to estimate disease-specific effects for all genes, while reducing uncertainty by borrowing confidence from the entire distribution of coefficients (internal shrinkage). Moreover, gene-specific effects help accounting for baseline differences across genes (codon usage, GC content, gene-specific mutation tendencies) which, if left unaccounted for, can skew modelling results. In this respect, instead of using variant counts, we could have used pNpS (polymorphic nonsynonymous-to-synonymous ratio). pNpS is similar to its non-polymorphic version, dNdS (non-polymorphic nonsynonymous-to-synonymous ratio), but less known. Metrics like pNpS and dNdS normalize for the actual number of nonsynonymous and synonymous opportunities in the sequence, however, ratio-based metrics have the disadvantage of being less interpretable in modelling. For these reasons, we decided to model gene-level nonsynonymous variant enrichment at polymorphic positions using variant counts, but then to use pNpS to confirm the model results. In fact, after modelling, we selected the genes that were categorized concordantly across CD and UC, for both Nsyn-decreased and Nsyn-increased genes, and then calculated their mean pNpS over subjects, for Control and IBD separately. Finally, we used these pNpS means to compute IBD-to-Control ratios and test statistical difference (Nsyn-decreased=1283, Nsyn-increased=43263). In the model, we also included dNdS, to approximate long-term genetic divergence existing between the strains within the subject and the reference genome, which could confound our results, when considering that different subjects could harbor strains that differ quite a lot in their genomic background. For this reason, we did not want to detect more nonsynonymous polymorphisms, only because some individuals might harbor a different uncharacterized subspecies.

### Functional enrichment

To retrieve gene’s KEGG Orthology (KO) terms and definitions we ran KofamScan on all the 20 genomes protein coding sequences provided by Prodigal, and selected only confident annotations (score > 0, N=361). Then, for each KO term we (1) extracted EC numbers from the KO terms definition, (2) retrieved KEGG pathway identifiers and definitions (from https://rest.kegg.jp/link/pathway/ko and https://rest.kegg.jp/list/pathway respectively), and (3) used the KEGGREST R package in batch mode to collect BRITE hierarchies. These last three steps allowed us to collect general functional annotation encompassing more KO terms together, while providing annotation for most of them. Before calculating odds-ratios we removed BRITE annotations reporting “Unknown function” or “General function prediction only” and excluded functions with less than 5 genes to reduce weakly supported results. Finally we manually curated annotations that reported eukaryotic-associated functions, opting for homologous annotations suitable for prokaryotes. Odds-ratios were calculated for each functional group by using raw frequency counts of genes as within one of the three gene evolutionary categories (*adaptive*, *neutrally-divergent* and *functionally-divergent*), based on their ND and Nsyn coefficients, with a smoothing correction (+0.5). In particular, odds-ratios were defined as follows = `cat_pos_in + 0.5)*(cat_neg_out + 0.5) / (cat_pos_out + 0.5)*(cat_neg_in + 0.5)`.

## Data availability

Scripts used for running the linear regression models and other analyses will be available at https://github.com/yassourlab/IBD_gene_diversity

